# A Novel Operant Conditioning Task to Assess Motivation to Exercise in Rats

**DOI:** 10.64898/2026.07.07.737013

**Authors:** Désirée R. Seib, Megan Q. Liu, Daniel J. Tobiansky, Stan B. Floresco, Kiran K. Soma

**Affiliations:** Department of Psychology and Djavad Mowafaghian Centre for Brain Health, University of British Columbia, Vancouver, BC, Canada; Department of Zoology, University of British Columbia, Vancouver, BC, Canada

**Keywords:** food restriction, brain, wheel running, mesocorticolimbic system, steroids

## Abstract

Voluntary physical activity is a highly motivated behavior with important implications for physical and mental health, yet the neural and endocrine mechanisms underlying motivation to exercise remain poorly understood. In contrast, motivation for sugar/palatable foods, drugs, and sex has been extensively characterized using operant paradigms. Here, we describe a novel progressive ratio operant task to measure motivation to run, independent of running ability. Using female Long Evans rats, which exhibit robust voluntary running behavior, we validated this paradigm by applying a manipulation well known to enhance the motivation to run: calorie restriction. Calorie-restricted animals exhibited increased operant responding to gain access to a running wheel, thus demonstrating heightened motivation for exercise. More specifically, calorie-restricted rats completed more ratios, reached a higher breakpoint in the progressive ratio task, ran more, and spent more time in the operant chamber. We did not observe any effects of calorie restriction on the estrous cycle or steroids (e.g. corticosterone, testosterone) in the blood or brain. Importantly, our task dissociates the motivational drive for physical activity from the ability to perform the physical activity itself, providing a new paradigm for studying the neural and endocrine mechanisms that regulate exercise motivation.

## Introduction

Exercise has been shown in numerous studies to enhance brain function, from neuropsychiatric disorders, such as depression and anxiety, to neurodegenerative diseases, such as Alzheimer’s and Parkinson’s disease (Umegaki et al., 2021; Zhou et al., 2017). Physical activity increases blood flow to the brain, promotes the survival of new neurons, and strengthens synaptic connections between mature neurons (Lehmann et al., 2020; Vonderwalde & Kovacs-Litman, 2018), which improve mood, memory, attention, and executive function (Lapmanee et al., 2017; Xiao et al., 2021). In contrast, sedentary behavior is associated with an increased risk of chronic diseases, cognitive decline, and mental health conditions (Pinto et al., 2023). Although it is well known that exercise is critical for the brain, cardiovascular system, and overall health, physical inactivity remains a global health concern, with about one-third of people leading sedentary lives (Park et al., 2020). Further research is needed to investigate the neural and hormonal mechanisms that drive motivation to exercise, which could lead to more effective strategies for increasing physical activity.

Neuroendocrine mechanisms play a crucial role in regulating physical activity, yet the motivation to exercise remains largely unexplored (Fragala et al., 2011). The brain’s regulation of the motivation to exercise is a complex interplay of factors, including energy balance, mood, and the perception of effort (Greenwood & Fleshner, 2019; Novak et al., 2012). Experimental and observational studies indicate that steroid hormones play a role in regulating physical activity. In male and female mice, gonadectomy reduces voluntary running (distance, duration, speed), while replacement with testosterone or estradiol reinstates activity levels (Bowen, 2012). Androgens and estrogens act not only on peripheral tissues such as muscle but also on central neural circuits that regulate motivation and activity (Jardí, Laurent, Dubois, et al., 2018; Lightfoot, 2008; Shay et al., 2026). In humans, epidemiological studies show complex correlations between habitual physical activity and androgen and estrogen levels, particularly in postmenopausal women (Bertone-Johnson et al., 2009; Hackney et al., 2024).

The mesocorticolimbic system regulates decision making, reward processing, and goal-directed behavior, including the motivation to pursue rewards (Seib et al., 2023). This system includes the ventral tegmental area (VTA) that sends dopaminergic projections to the nucleus accumbens (NAc) and the medial prefrontal cortex (mPFC). Dopamine signaling regulates motivation for sucrose rewards and mating behavior (Balfour et al., 2004). With respect to dopamine and physical activity, long-term (6 weeks) exercise in rats induces changes in the mesocorticolimbic system, such as increased activity in the NAc and increased tyrosine hydroxylase (TH) mRNA expression in the VTA (Greenwood et al., 2011). However, little is known about the neural mechanisms that drive motivation for exercise. Inactivation of the NAc or the mPFC in rats reduces the motivation to run (Basso & Morrell, 2015). Despite its well-established role in regulating motivation, a significant gap remains in understanding how the mesocorticolimbic system regulates motivation to engage in running behavior.

Steroid hormones, including glucocorticoids (e.g., corticosterone) and sex hormones (e.g., testosterone, estradiol), modulate the mesocorticolimbic system (Goel et al., 2014; Seib et al., 2023; Tobiansky, Wallin-Miller, et al., 2018). Corticosterone influences energy balance and responses to stress, which can either enhance or suppress exercise motivation, depending on stress intensity and duration. Chronic stress reduces running in mice (DeVallance et al., 2017), while an acute increase in corticosterone increases running (Duclos et al., 2009; Ebada et al., 2016). Sex steroids can enhance running by acting on dopaminergic pathways in the mesocorticolimbic system. Androgens increase physical activity and promote reward sensitivity (Jardí, Laurent, Dubois, et al., 2018; Jardí, Laurent, Kim, et al., 2018). Estrogens modulate mood and cognitive function, potentially enhancing the perceived reward of physical activity and promoting exercise (Anantharaman-Barr & Decombaz, 1989; Dreher et al., 2007; Kokane & Perrotti, 2020; Krause et al., 2021; Satta et al., 2018).

Belke and colleagues have studied the motivation to run by using running as a reward in operant conditioning paradigms with rodents. Temporal parameters, such as the immediacy of wheel access or the duration of the reward, influence preference and response rates (Belke, 1997; Belke & Wagner, 2005). Moreover, dopaminergic agents, such as amphetamine, reduce the motivation to run (Belke et al., 2019). These findings highlight how both the timing of rewards and neurochemical state modulate the motivational significance of physical activity. In addition, Cordony and colleagues used a progressive ratio (PR) task (a measure of “wanting”), in which the required number of presses increases after each earned reward. This study demonstrated that restricting access to a running wheel increases the motivation to regain access; however, a greater amount of prior running experience did not affect this motivation (Cordony et al., 2019). Our study builds on this work by employing a more complex operant conditioning system that enables the examination of specific components of behavioral mechanisms in greater detail.

Here, our goal was to establish a novel behavioral task that could separate two aspects of exercise: the amount of running versus the motivation to run, using an automated operant conditioning procedure. Previous research shows that calorie restriction increases the motivation to run (Belke et al., 2016, 2017; Belke & Pierce, 2016). We used this established relationship to test our newly developed task, aiming to provide a more nuanced understanding of exercise behavior. Using highly specific, sensitive, and precise liquid chromatography tandem mass spectrometry, we also investigated whether calorie restriction altered steroid levels in the blood and mesocorticolimbic system of female rats.

## Methods

### Animals

All procedures were approved by the University of British Columbia Animal Care Committee and were in accordance with the guidelines of the Canadian Council on Animal Care. Adult female Long Evans rats (60 to 68 days old, n=23, Charles River Laboratories) were housed in a temperature-controlled vivarium (22°C; 40 to 50% relative humidity) on a reversed 12 h light:12 h dark cycle (lights on from 19:30 h to 07:30 h) in the UBC Centre for Disease Modeling. We used female rats because they are more motivated to run than male rats (Eikelboom & Mills, 1988; Tanner et al., 2022). Upon arrival and before calorie restriction, all rats were given *ad libitum* access to a standard rodent chow (Teklad Diet 2918; Envigo, Indianapolis, IN, USA). Throughout the experiment, all rats had free access to drinking water (purified by reverse osmosis and sterilized by chlorination). The rats were pair-housed in Allentown cages and left undisturbed for one week to acclimate to the colony room. Each cage was supplied with beta chip bedding as well as a PVC pipe and paper towels for enrichment.

Following acclimation, the rats were group-housed (four rats per cage) for two weeks in Rubbermaid Tupperware containers (36.8 cm × 68.5 cm × 50.8 cm), and each container had one running wheel. During this period, rats had free access to the running wheel for acclimation. After wheel acclimation, rats were once again pair-housed in Allentown cages without a running wheel. For the next nine days, all rats were handled daily.

### Calorie restriction

Upon their return to pair-housing, all rats were weighed daily, and each cage was randomly assigned to either the calorie restricted (CR) or control (CON) group (n=11-12/group). Rats in each group were housed together; no CR rat was pair-housed with a CON rat. Food hoppers of CON cages were topped up daily. In a previous study investigating the motivation to run using operant conditioning, rats that were progressively calorie restricted displayed peak wheel running and motivation when they reached 75% of their initial weight (Pierce et al., 1986). Thus, in our study, the average weight of CR rats was restricted to 80% of that of CON rats throughout the experiment. CR rats were initially fed 2% of their body weight in chow each day. Once the average CR weight reached 80% of the average CON weight, CR rats were fed an amount of chow weighing 10% of their body weight (approximately 14 g) each day. Body weights were monitored closely, and the amount of chow given daily to each CR rat was adjusted carefully. Each CR rat was fed once per day immediately after operant training or testing, in an individual cage to ensure each rat received the appropriate amount of chow. Rats were given nine days to habituate to calorie restriction before beginning operant training.

### Apparatus

Operant training and testing sessions were conducted in operant chambers (20.3 cm × 16 cm × 17.8 cm; Med Associates Inc., St. Albans, VT, USA). Each operant chamber was attached to a running wheel (36 cm × 10 cm, 1.13 m circumference; Model ENV-042A, Med Associates Inc.) via a guillotine door. The running wheels were each equipped with an electronic brake (Model ENV-042A-SBA, Med Associates Inc.). Each operant chamber was in a sound-attenuating box that included a fan for ventilation and a small window to monitor the rat’s activity. Two retractable levers were on one wall of the main chamber, and a cue light was above each lever. Only the left lever and cue light were used in this study. A house light was above the cue lights. Behavioral data were recorded with Med-PC (Med Associates Inc.).

### Operant training

All operant training and testing sessions were conducted within the first 3 h of the dark period, when rats are most active (Eikelboom & Mills, 1988). The first day of operant training was also the first day all subjects underwent a daily vaginal lavage for estrous cycle staging (described below).

The first stage of operant training was wheel training and began nine days after the start of calorie restriction. This stage familiarized the rats with the operant chamber and attached wheel. During wheel training, the house light was on, the guillotine door was open, and the running wheel brake was disengaged for the entire session. Moreover, both levers were retracted, and the cue lights remained off throughout the session. Each session lasted 30 min, during which the number of wheel rotations was recorded. Once the subject reached the criterion of 500 rotations in a session, they were advanced to the next stage of operant training. All subjects met this criterion during their first wheel-training session.

The second stage of operant training taught the rats to associate lever pressing with gaining access to the running wheel by using a fixed ratio (FR) schedule of reinforcement. Each lever training session lasted 30 min, and each subject had one session per day. All subjects began with FR1, in which one lever press was required to gain access to the unlocked wheel. All FR training sessions began with both levers retracted, all lights off, and the guillotine door shut. After 1 min, the left lever (closest to the guillotine door) was extended, and its cue light was turned on. Once a subject pressed the lever to obtain a reward, the lever was retracted, its cue light was turned off, the house light was turned on, the guillotine door was opened, and the brake was disengaged. The subject could run in the wheel for 1 min before the brake was reengaged. Then the house light was turned off, the lever extended, and the cue light turned on. The guillotine door remained open for the remainder of the session. The subject then had to return to the main chamber and lever press again to disengage the brake. Subjects were given 30 min to reach the criterion of achieving 10 rewards (i.e. access to the wheel). When a rat reached the criterion of 10 rewards under FR1, it was moved on to a FR2 schedule. Once 10 rewards were reached within one session under FR2, subjects were advanced to FR5, where daily training sessions continued until an animal again met the criterion of 10 rewards in a session. In the following session, a rat was advanced to a progressive ratio (PR) task (described below). Three subjects (two CON animals and one CR animal) did not reach 10 rewards in one session under FR5 but were advanced to the PR task due to time constraints; all individuals successfully performed the PR task. Subjects spent a maximum of 14 days in FR training.

### Progressive ratio task

This operant task was used to assess the rats’ motivation to use a running wheel. Under a PR schedule of reinforcement, the number of lever presses required to gain access to the wheel increased after each ratio was completed. Our PR schedule increased in a series derived from the following equation (Richardson & Roberts, 1996):

*Lever press requirement (rounded to nearest integer)* = 5*e*^0.2 ⁢ *trail number*^ − 5

The resulting series was: 1, 2, 4, 6, 9, 12, 15, 20, 25, 32, 40, 50, 62, 77, 95, 118, 145, 178. The rat initially pressed the lever once to gain access to the wheel for 1 min. To regain access, the rat had to press the lever twice, and then four times, and so on.

Each PR session began with all lights off in the operant chamber, both levers retracted, the guillotine door closed, and the wheel brake engaged. After 1 min, the left lever was extended, and its cue light was turned on. Rats were given 10 min to reach the ratio requirement. If the criterion was reached, the lever was retracted, its cue light was turned off, the house light was turned on, the door was opened, and the brake was disengaged. The rats could run in the wheel for 1 min. After 1 min, the brake was reengaged, the house light was turned off, the left lever was extended, and its cue light was turned on. The guillotine door remained open for the remainder of the session. The rat needed to meet the next criterion within 10 min to regain access to the unlocked wheel. If the rat failed to reach the next criterion within 10 min, then that criterion was recorded as their breakpoint (i.e., the criterion not completed) and the session was then terminated. The number of wheel rotations, number of times access to the wheel was obtained (i.e., rewards obtained), breakpoint, latency to complete each ratio, and number of pauses in lever pressing were recorded. A pause in lever pressing was defined as any period of lever press inactivity greater than 5 sec while the wheel was locked and the lever was extended.

All subjects underwent the PR task once daily for 24 days. The first 10 days were considered the PR training phase, and the last 14 days were considered the PR testing phase. One subject in the CR group died for unknown reasons on day 4 of PR training (resulting in n=11), so all data from that subject were excluded. After each session, CR subjects were immediately fed before being returned to their home cage. The Med-PC code for this study is available by emailing the corresponding author.

### Estrous cycle staging

Immediately after each operant training or testing session, vaginal lavages were performed as before (Seib et al., 2025). Daily vaginal lavage began on the first day of operant training. Subjects were classified as proestrus, estrus, or diestrus based on the cellular morphology and density (Cora et al., 2015). Given the short length of metestrus (6-8 h), the longer length of diestrus (48-72 h), and the subtle differences in cell morphology between these two stages, any smear that resembled either metestrus or diestrus was classified as diestrus (Ajayi & Akhigbe, 2020).

### Tissue collection

All subjects were euthanized the day after the last PR testing session between 09:30 and 10:30 h (2-3 h after lights off). Animals in the CR group were last fed after the last PR session. Subjects were rapidly and deeply anesthetized with isoflurane before rapid decapitation. Euthanasia was completed within 3 min of initial cage disturbance to avoid stress-induced changes in steroids (Taves et al., 2011). Trunk blood was collected in a microcentrifuge tube and flash-frozen on dry ice. The brain was rapidly extracted and flash-frozen on powdered dry ice. Blood and brain samples were stored at -70°C.

### Brain microdissection

Brains were sectioned at 300 μm in the coronal plane using a cryostat and mounted onto glass Superfrost Plus slides, as before (Tomm et al., 2022). Brain sections were stored at -70°C. Sections were microdissected on dry ice using the Palkovits punch technique with a punch tool (1-mm diameter; 0.244 mg/punch; Thermo Fisher Scientific, Waltham, MA, USA) (Figure S1). Punches were collected from regions in the mesocorticolimbic system and associated regions: mPFC, NAc, bed nucleus of the stria terminalis (BNST), amygdala (AMY), dorsal hippocampus (dHPC), ventral hippocampus (vHPC), and VTA. Punches were also collected from the preoptic area (POA) and hypothalamus (HYP), which regulate running and food intake (De Rijke et al., 2005). Regions were identified using major anatomical landmarks and an adult rat brain atlas (Paxinos & Watson, 2007). Punches were collected bilaterally (4 to 8 punches per region). Punches were placed into cold 2 mL polypropylene bead ruptor tubes (Sarstedt AG & Co., Nümbrecht, Germany) and stored at - 70°C.

### Steroid extraction

All samples were processed using liquid-liquid extraction as before (Hamden et al., 2021; Jalabert et al., 2021; Tobiansky, Korol, et al., 2018). In brief, five zirconium ceramic beads (1.4-mm diameter) were added to each bead ruptor tube. To track steroid recovery in each sample, we added 50 μL of deuterated internal standards (corticosterone-d8, progesterone-d9, E17β-estradiol-d4, and testosterone-d5) in 50:50 high-performance liquid chromatography (HPLC)-grade methanol:MilliQ water. Then 1 mL of HPLC-grade acetonitrile (ACN) was added, and samples were homogenized at 4 m/s for 30 sec using a bead mill homogenizer. Next, samples were centrifuged at 16,100 *g* for 5 min, and 1 mL of supernatant was transferred to pre-cleaned borosilicate glass culture tubes. Then 0.5 mL of HPLC-grade hexane was added, and samples were vortexed and centrifuged at 3,200 *g* for 2 min. The hexane was discarded, and the ACN was dried in a vacuum centrifuge at 60°C for 45 min. Samples were resuspended in 55 μL of 25% HPLC-grade methanol in MilliQ water, transferred to 0.6 mL polypropylene microcentrifuge tubes, and centrifuged at 16,100 *g* for 2 min. Then 50 μL of supernatant were transferred to a LC vial insert, and samples were stored at -20°C until analysis by liquid chromatography tandem mass spectrometry (LC-MS/MS). Samples were extracted alongside calibration curves (0.05 to 1000 pg per tube), blanks, and quality controls (QCs).

### Steroid analysis by LC-MS/MS

Six steroids were measured in blood (20 μL) and brain tissue (0.976-1.952 mg): corticosterone, 11-dehydrocorticosterone (DHC), progesterone, 17β-estradiol, testosterone, and androstenedione. Steroids were quantified using a QTRAP 6500 UHPLC-MS/MS system (Sciex LLC, Framingham, MA, USA) as before (Jalabert et al., 2021). In brief, 45 μL was injected into a Shimadzu Nexera X2 UHPLC system, passed through an in-line filter and guard column, and separated on a Kinetex® 2.6 µM EVO C18 100 A° LC column (2.1 × 50 mm; 2.6 μm; at 40°C) using 0.1 mM ammonium fluoride in MilliQ water as mobile phase A (MPA) and HPLC-grade methanol as mobile phase B (MPB) (Jalabert et al., 2024). We used electrospray ionization (positive and negative modes). We used multiple reaction monitoring (MRM), with two MRM transitions for analytes and one MRM transition for internal standards.

### Data analysis

A two-way mixed ANOVA was used to analyze body mass. Group (CON and CR) was the between-subjects factor, and week (0 to 6, 0 = start of CR) was the within-subjects factor.

Two-way mixed ANOVAs were also used to analyze behavioral data over the 14-day PR testing phase. Separate ANOVAs were conducted on the dependent variables: completed ratios, breakpoint, total rotations, rotations per ratio, total time, total pauses, and pauses per ratio. For all analyses, treatment group (CON and CR) was the between-subjects factor, and PR session (time) was the within-subjects factor. For behavioral data averaged over the PR testing phase, two-tailed independent t-tests were used. Data distributions were assessed for normality using the Shapiro-Wilk test. If the assumption of normality was violated, we performed the analysis on log-transformed data.

Steroids were considered below the lower limit of quantification (LLOQ) if they fell below the lowest standard on the calibration curves. For groups with ≥ 20% samples above the LLOQ for a given analyte, missing values were imputed via left-censored quantile regression imputation using the MetImp web tool (Hamden et al., 2021; Jalabert et al., 2021; Wei et al., 2018). Independent samples t-tests were used to compare blood steroid levels between the CON and CR groups. Two-way mixed ANOVAs were used to compare brain steroid levels, with group (CON and CR) as the between-subjects factor and region (mPFC, NAc, BNST, POA, AMY, HYP, dHPC, vHPC, VTA) as the within-subjects factor.

For all analyses, α was set at ≤ 0.05. All statistics were analyzed, and all figures were generated using GraphPad Prism version 10 (GraphPad Software Inc., USA). The data presented in all figures are non-transformed (mean ± SEM).

## Results

### CR and body mass

Initially, female rats were housed in a large cage with a running wheel in groups of four for two weeks. Then, the body mass of CR rats was reduced to and maintained at 80% of *ad libitum*-fed CON rats throughout the experiment, to examine the effect of calorie restriction on the motivation to run (Figure 1; group: F_1,21_=62.83; *P*<0.0001; time: F_1.42,29.86_=95.25; *P*<0.0001; group x time: F_1.42,29.86_=59.05; *P*<0.0001). After the CR group reached 80% of the weight of the CON group, operant training began by allowing rats to run in the wheel attached to the operant chamber for 30 min. Subsequently, rats were trained to lever press using FR training, followed by training on the PR task. Once behavior stabilized, rats were tested for 14 consecutive days on the PR task (Figure 2).

**Figure 1.**
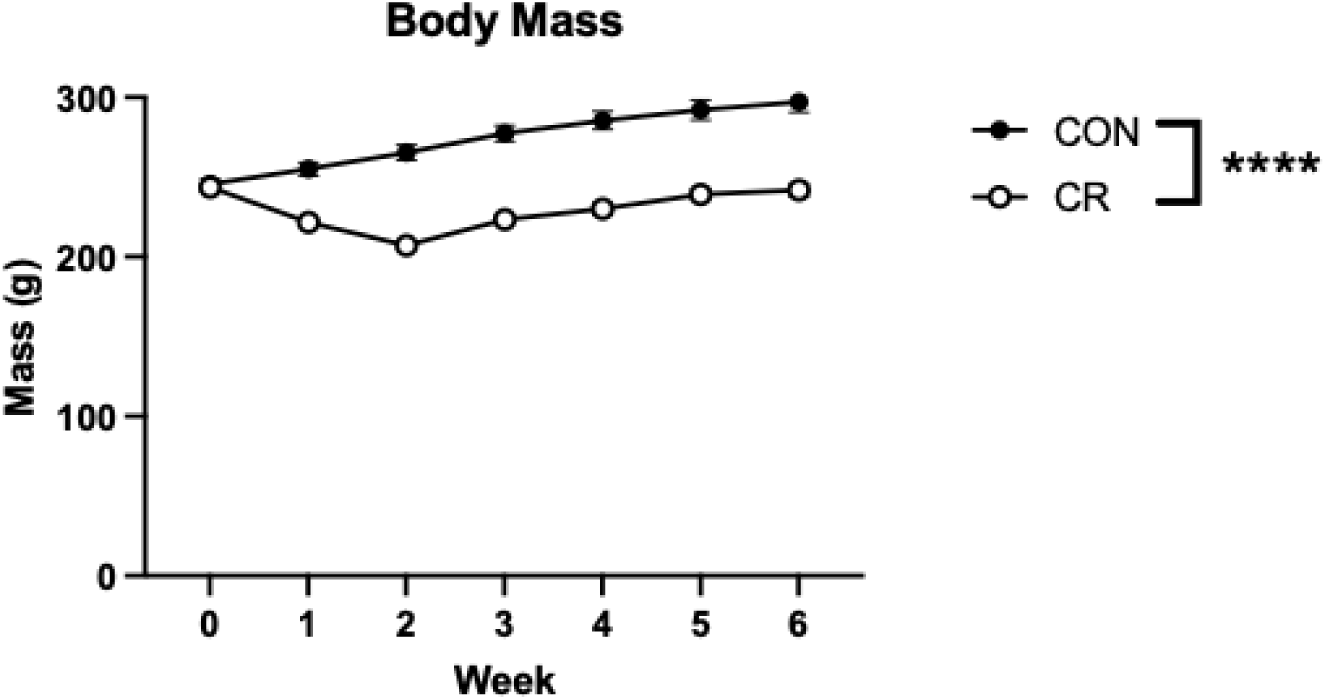
Body mass. Body mass of control (CON) and calorie-restricted (CR) female rats. The average weight of CR rats was maintained at 80% of the average weight of CON rats for the duration of the experiment. ****p<0.0001.

**Figure 2.**
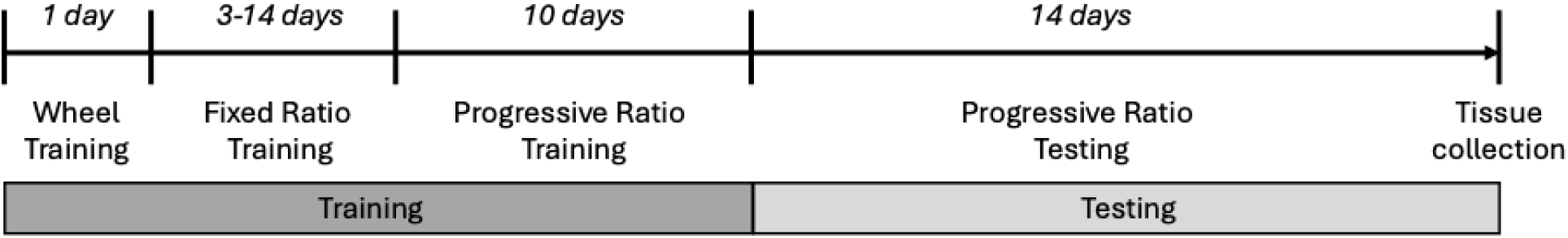
Experimental timeline. After the start of calorie restriction for the calorie-restricted (CR) group, both the control (CON) and CR groups of female rats completed wheel training and operant training, followed by 14 days of progressive ratio testing.

### CR increases the motivation to run of female rats in the PR operant task

Completed ratios were measured as a direct assessment of motivation and to quantify task engagement and effort expenditure, reflecting the rats’ capacity and willingness to perform the required presses to obtain access to the running wheel. CR female rats completed significantly more ratios compared to CON rats (Figure 3A; group: F_1,21_=4.593; *P*=0.044; time: F_6.76,141.90_=0.790; *P*=0.593; group x time: F_6.76,141.90_=0.472; *P*=0.848 and Figure 3B; *t*_21_=2.143; *P*=0.044).

**Figure 3.**
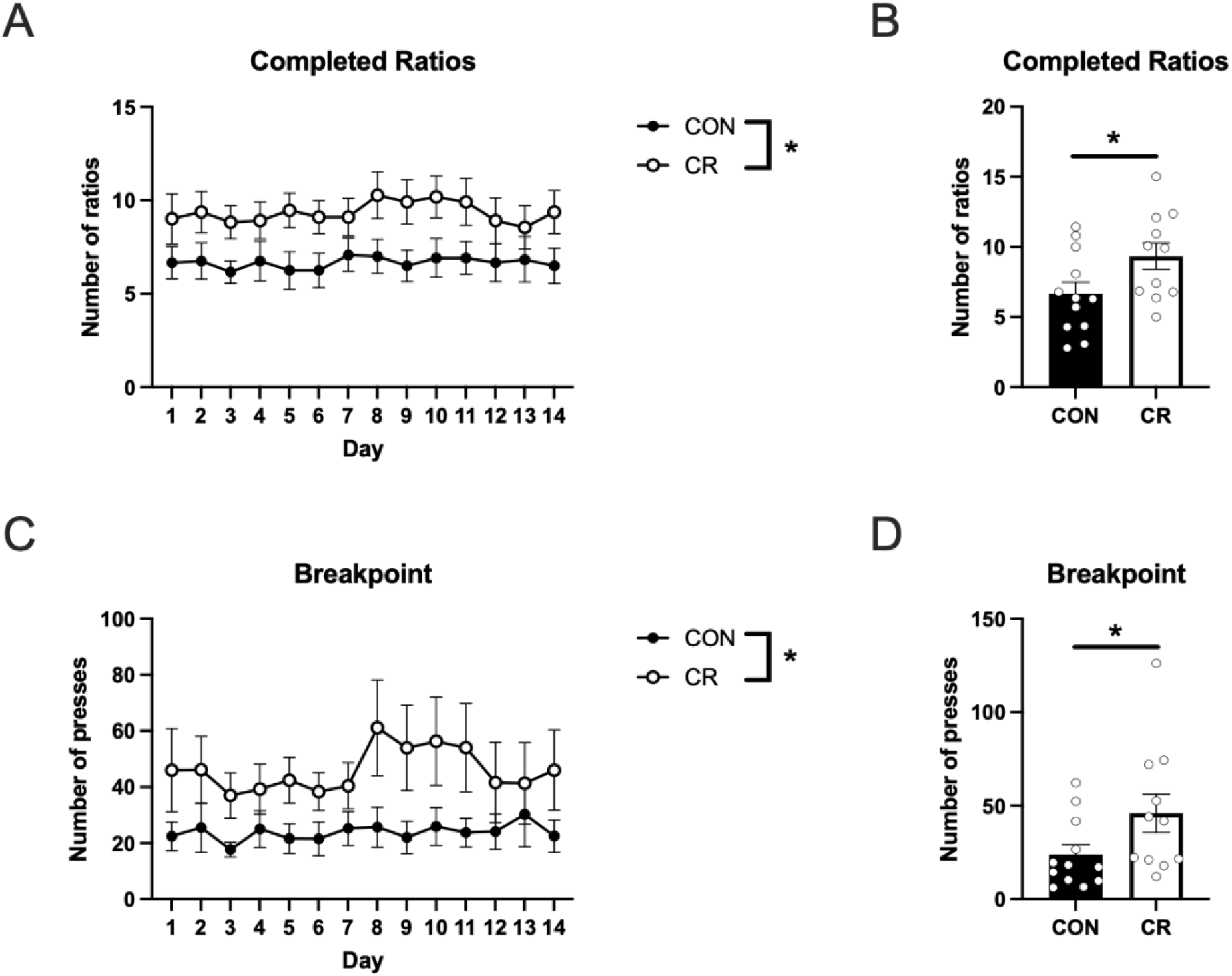
Motivation to run. (A) Calorie-restricted (CR) females completed significantly more ratios per session than control (CON) females. (B) CR females completed significantly more ratios per session than CON females, when individual subjects were averaged over the 14-day testing period. (C) CR females had a significantly higher breakpoint than CON females. (D) CR females had a significantly higher breakpoint than CON females, when individual animals were averaged over the 14-day testing period. *p<0.05.

Breakpoint was measured as an index of motivation, defined as the response requirement not completed and the point of task disengagement, reflecting the limit of effort rats were willing to exert to access the running wheel. CR significantly increased the breakpoint (Figure 3C; group: F_1,21_=4.793; *P*=0.040; time: F_7.39,155.20_=0.855; *P*=0.549; group x time: F_7.39,155.20_=0.436; *P*=0.887 and Figure 3D; *t*_21_=2.212; *P*=0.038).

### CR increases the running activity of female rats in the PR operant task

In line with the observed increase in completed ratios, CR rats ran more than CON rats, as measured by the number of total rotations during a session (Figure 4A; group: F_1,21_=9.76; *P*=0.005; time: F_6.03,126.60_=0.995; *P*=0.432; group x time: F_6.03,126.60_=0.427; *P*=0.861) and throughout the 14 test days (Figure 4B; *t*_21_=3.125; *P*=0.005). In addition, the increase in the motivation or ability to run was also apparent as CR rats completed more rotations per ratio (Figure 4C; group: F_1,21_=15.49; *P*<0.001; time: F_6.36,131.20_=1.025; *P*=0.414; group x time: F_6.36,131.20_=0.461; *P*=0.846 and Figure 4D; *t*_21_=3.935; *P*<0.001).

**Figure 4.**
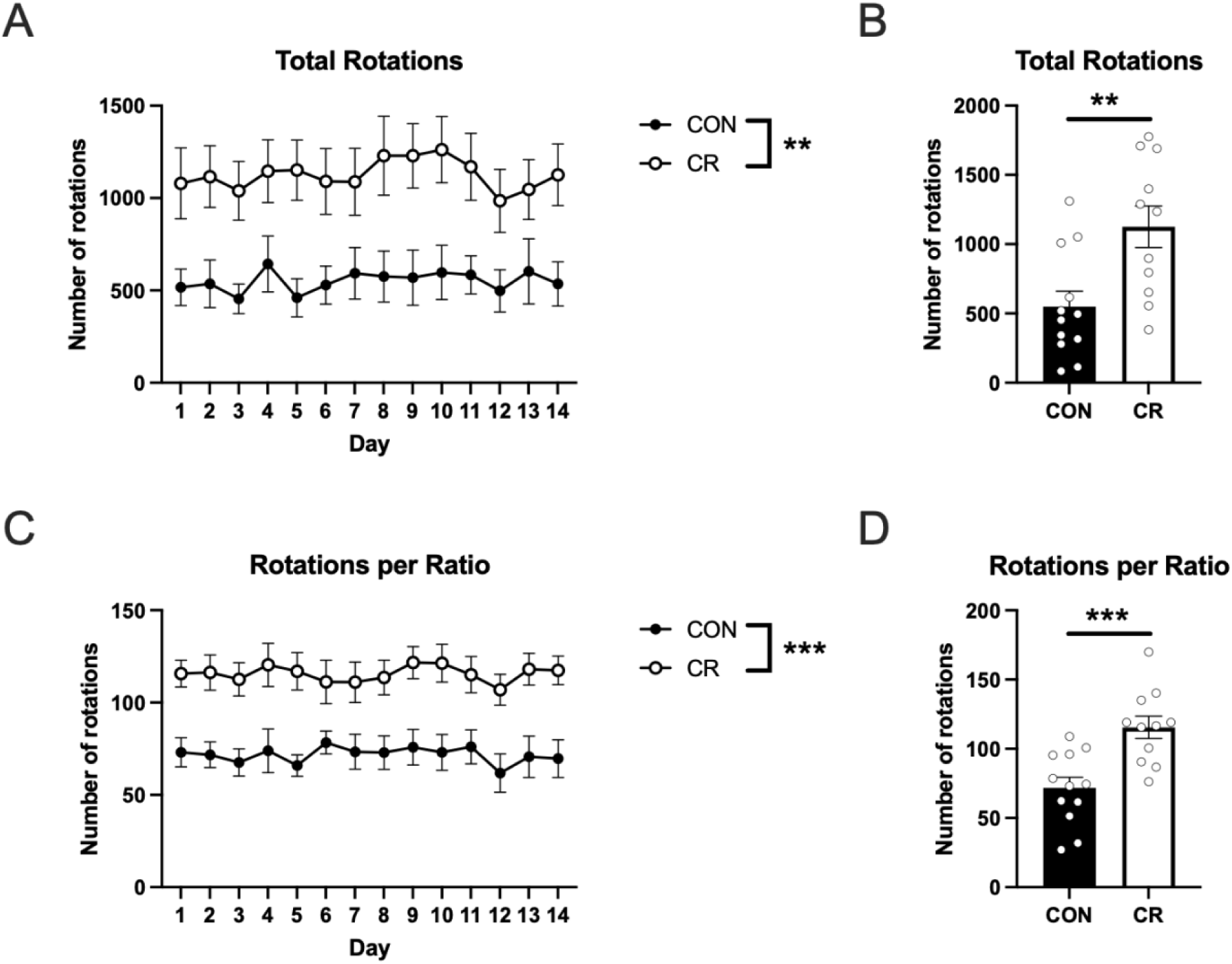
Running activity. (A) Calorie-restricted (CR) females completed significantly more total rotations per session than control (CON) females. (B) CR females completed significantly more total rotations per session than CON females, when individual subjects were averaged over the 14-day testing period. (C) CR females completed significantly more rotations per ratio than CON females. (D) CR females completed significantly more rotations per ratio than CON females, when individual subjects were averaged over the 14-day testing period. **p<0.01; ***p<0.001.

### Effects of CR on temporal and operational parameters

Total time spent in the operant chamber per session was significantly increased by CR (Figure 5A; group: F_1,21_=7.547; *P*=0.012; time: F_6.90,144.90_=0.859; *P*=0.540; group x time: F_6.90,144.90_=1.195; *P*=0.310 and Figure 5B; *t*_21_=2.747; *P*=0.012). In addition, the total number of pauses per session was significantly increased in CR rats (Figure 5C; group: F_1,21_=8.061; *P*<0.01; time: F_6.82,143.20_=0.941; *P*=0.476; group x time: F_6.82,143.20_=1.426; *P*=0.201 and Figure 5D; *t*_21_=2.839; *P*=0.010). However, the frequency of pauses per completed ratio was not significantly altered by CR (Figure 5E; group: F_1,21_=1.787; *P*=0.196; time: F_7.85,161.90_=1.006; *P*=0.433; group x time: F_7.85,161.90_=1.970; *P*=0.055 and Figure 5F; *t*_21_=1.282; *P*=0.214). This indicates that the increase in the total number of pauses in the CR group is attributable to these animals achieving higher ratios over the session. Collectively, these findings suggest that the total number of pauses increased as a function of time spent in the operant chamber, and did not reflect an increase in behavioral disengagement due to CR.

**Figure 5.**
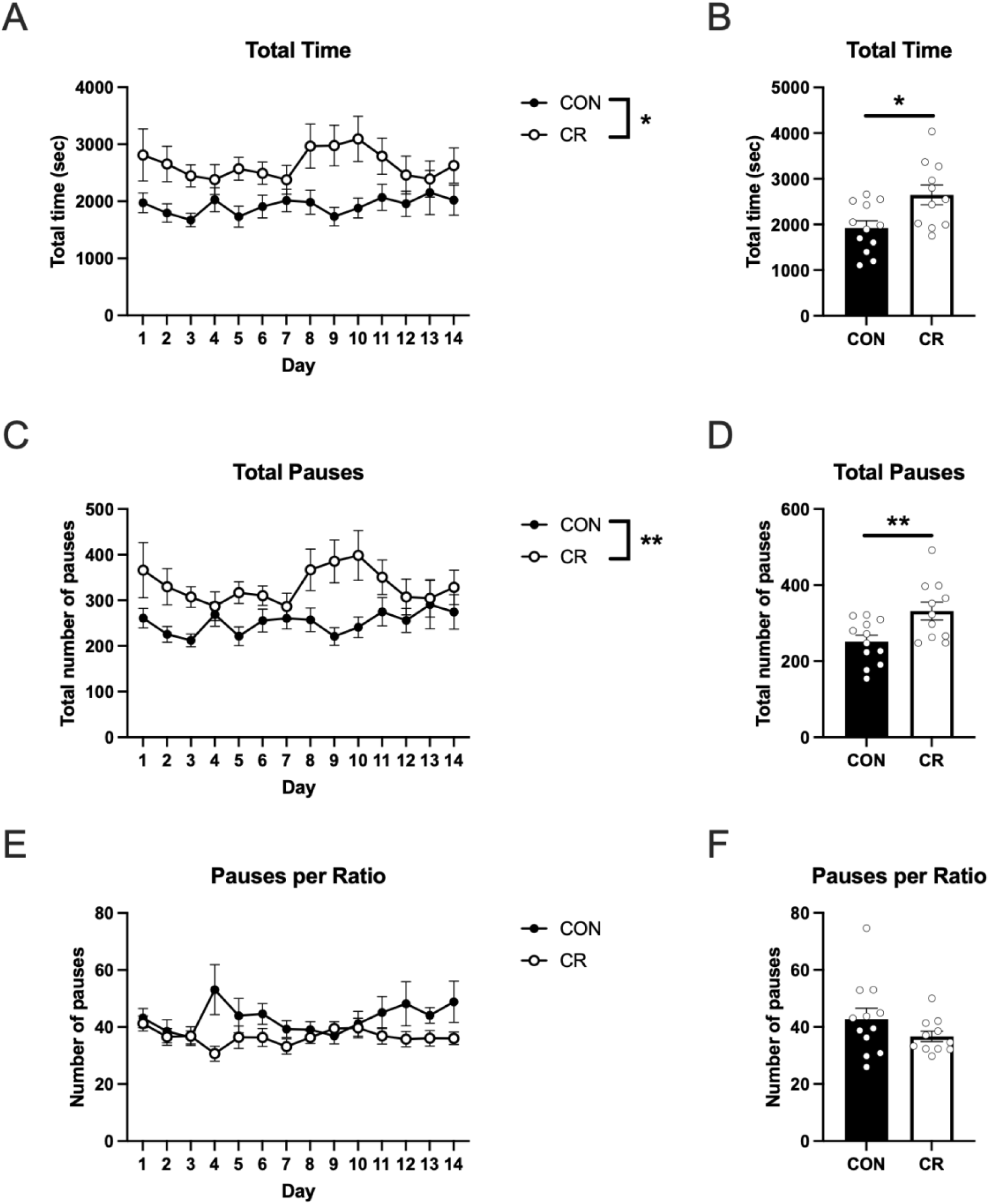
Temporal and operational parameters. (A) Calorie-restricted (CR) females spent significantly more time in the operant chambers (performing the PR task) per session than control (CON) females. (B) CR females spent significantly more time in the operant chambers per session than CON females, when individual subjects were averaged over the 14-day testing period. (C) CR females made significantly more total pauses per session than CON females. (D) CR females made significantly more total pauses per session than CON females, when individual subjects were averaged over the 14-day testing period. (E) CR and CON females did not significantly differ in the number of pauses per ratio during an individual session or (F) when averaged over the 14-day testing period. *p<0.05: **p<0.01.

### CR did not alter estrous cycling and steroid hormones in female rats

CR females showed no changes to their estrous cycle, indicating that mild food restriction did not affect the estrous cycle in Long-Evans female rats (data not shown). Additionally, the estrous stage did not impact performance in the PR task (data not shown).

After PR testing, we measured levels of six steroids (corticosterone, DHC, progesterone, estradiol, testosterone, and androstenedione) in behaviorally relevant brain regions (mPFC, NAc, BNST, POA, HYP, AMY, dHPC, vHPC, and VTA) and in the blood. Estradiol levels were too low to be reliably detected in most samples. Corticosterone, DHC, and progesterone (Table S1) and the androgens testosterone and androstenedione (Table S2) in the brain and blood were not significantly altered by CR.

## Discussion

We developed a novel and robust behavioral paradigm to measure the motivation for physical activity in rats. Our results show that calorie restriction in female Long Evans rats to 80% of the weight of an *ad libitum*-fed CON group increases wheel running activity, as CR rats completed more ratios and reached a higher breakpoint in the progressive ratio task. In addition, CR rats showed increased total rotations and rotations per ratio compared to the CON group. Thus, CR increased the motivation to run, as expected from previous work (Belke & Pierce, 2016). This behavioral paradigm will be very useful for examining the environmental, neural, and endocrine factors that impact the motivation for exercise.

### Key features of the behavioral paradigm

This research addresses a major knowledge gap in understanding the neuroendocrine regulation of the motivation for exercise. The neuroendocrine bases of motivation for sugar rewards and sexual behavior have been studied in great detail (Balfour et al., 2004; Dokovna et al., 2019; Floresco et al., 2008; Ghods-Sharifi & Floresco, 2010), while the neural and hormonal influences on motivation for exercise remain basically understudied. One challenge in studying physical activity is distinguishing between the *motivation* for physical activity and the *ability* to perform physical activity (i.e., wanting access to a running wheel versus the number of rotations) (Careau et al., 2026).

In this study, we present a novel paradigm using commercially available equipment that can be employed to measure running and the motivation to run. We selected this experimental setup based on several considerations. Animals were readily trained on the task, as two weeks of pre-exposure to a running wheel ensured that rats learned to run consistently (Belke & Pierce, 2016). Female Long-Evans rats were chosen because mild food restriction does not disrupt the estrous cycle of Long-Evans rats (Tropp & Markus, 2001), their running activity is not influenced by estrous stage if running periods are short (Belke & Pierce, 2016), and females show higher motivation to run than males in this and other rat strains (Mathis et al., 2024; Tanner et al., 2022). Additionally, Long-Evans rats display greater running motivation than other commonly used strains such as Sprague-Dawley or Wistar rats (Bauer, 1990). Our preliminary data suggest that male Long-Evans rats are also easily trained to perform this task (Rutledge, 2021). Last, all testing was conducted during the early dark phase, when running activity is maximal (Eikelboom & Mills, 1988).

### Calorie restriction as a proof of principle

Our results support the validity of the present apparatus and novel program for assessing motivation to run by replicating key findings reported by Belke and colleagues, who demonstrated that mild food restriction increases motivation to access a running wheel (Belke & Pierce, 2016). Using a similar paradigm, animals maintained at 90% of their free-feeding body weight exhibited greater motivation to obtain wheel access, without evidence of physiological stress. Here, an 80% calorie restriction is sufficient to elevate motivation for running but remaining within a range that does not induce a stress response. Indeed, higher food restriction elevates stress markers when imposed as a severe or forced manipulation (Han et al., 2001; McGhee et al., 2009), whereas milder or voluntary restriction produces fewer stress-related effects (Finnell & Ferrario, 2025), indicating that the degree of food restriction is a critical determinant. Here, calorie restriction increased running performance, as reflected by a greater number of wheel rotations per session and more rotations per completed ratio during the 1 min wheel access period.

CR animals also displayed higher operant responding, completing more ratios across the two-week testing period and achieving a higher breakpoint. Moreover, CR animals spent more total time in the operant chamber and had a higher absolute number of pauses. However, when pauses were normalized to the number of ratios completed, CR animals did not exhibit more pauses per ratio, indicating that the increase in pauses was simply a consequence of their greater overall task engagement (i.e., time spent in the operant chamber). Thus, CR did not just increase the amount of running but also increased the motivation to work for access to running, similar to previous studies (Belke & Pierce, 2016; Pierce et al., 1986).

The findings of Belke and colleagues are relevant for interpreting the effects of food restriction in the present study, as their work demonstrates that restricted feeding alters the motivational value of running rather than simply the rats’ physical ability to run (Belke et al., 2017). Food-restricted rats work harder to obtain access to a running wheel, particularly under high intensity schedules, indicating that metabolic state alters the reinforcing value of exercise and sets the stage for potential dissociations between motivation to obtain access to the wheel and the amount of running performed (Belke & Pierce, 2016). This distinction suggests that elevated running under food restriction reflects increased incentive value of wheel access, not merely heightened activity levels.

We did not observe any changes in steroid levels in the blood or brains of female rats following calorie restriction. Thus, the observed changes in the motivation to run might not be driven by changes in steroid levels. Potential mechanisms driving the motivation to run could include non-steroidal hormone pathways such as ghrelin signaling (Clifford et al., 2011; Mifune et al., 2020). Furthermore, although food restriction can alter glucocorticoid levels (Han et al., 2001), the mild food restriction used in this study did not elicit a corticosterone response.

While it may seem counterintuitive that calorie restriction prompts an increase in physical activity, this behavioral shift reflects a conserved evolutionary survival mechanism. During long-term energy deficits, ghrelin signaling overrides energy conservation cues to activate the mesocorticolimbic pathway (Mifune et al., 2020). This shift prevents lethargy and instead induces hyperlocomotion, often manifested as an acceleration in spontaneous wheel running, thereby compelling the organism to seek novel food sources (Novak et al., 2012).

Together, these findings highlight the importance of separating measures of motivation from measures of performance when examining physical activity. This paradigm may serve as a framework for future studies aimed at examining the neural mechanisms underlying the motivation to run with greater precision and detail.

### Future directions

Our approach may be useful for examining how pharmacological treatments influence the motivation to exercise, an issue that remains poorly understood. Many widely used therapies, including cancer treatments such as abiraterone, have negative metabolic side effects (Decamps et al., 2023; Kiwata et al., 2016). It is unclear whether these metabolic effects arise directly from the drug or indirectly through reductions in physical activity. The present behavioral paradigm could be used to assess how pharmacological treatments, including dopaminergic drugs, antiandrogens, and estrogen receptor antagonists alter the motivation to run. Further, region-specific inactivation studies can identify the neural circuits involved (Brown et al., 2014). Our findings and novel task can also help future studies parse out how and why calorie restriction enhances running motivation, potentially reflecting an evolutionarily conserved drive to seek food during an energy deficit. More broadly, this work may identify strategies to enhance motivation for exercise, an alternative to “exercise mimetics,” thereby promoting physical activity and reducing the health risks associated of inactivity.

## Acknowledgements

This work was supported by a Discovery Grant from the Natural Sciences and Engineering Research Council of Canada (NSERC) to SBF (RGPIN-2024-04207), a Project Grant from the Canadian Institutes of Health Research (CIHR) to KKS and SBF (169203), a postdoctoral fellowship from the Social Exposome Cluster and Human Early Learning Partnership (HELP) of the University of British Columbia to DRS, the University of Prince Edward Island Jeanne and J. Louis Lévesque Research Chair in Human Health to DRS, a CIHR CGSM to MQL, and Bluma Tishler Award to DJT. We thank Griffin Rutledge, Whitney Krieger, Valerie Lo, Hui W. Chen, Anastasia Korol, and Asmita Poudel for help with experimental procedures.

