## Supplemental Material for "A Novel Operant Conditioning Task to Assess Motivation to Exercise in Rats"

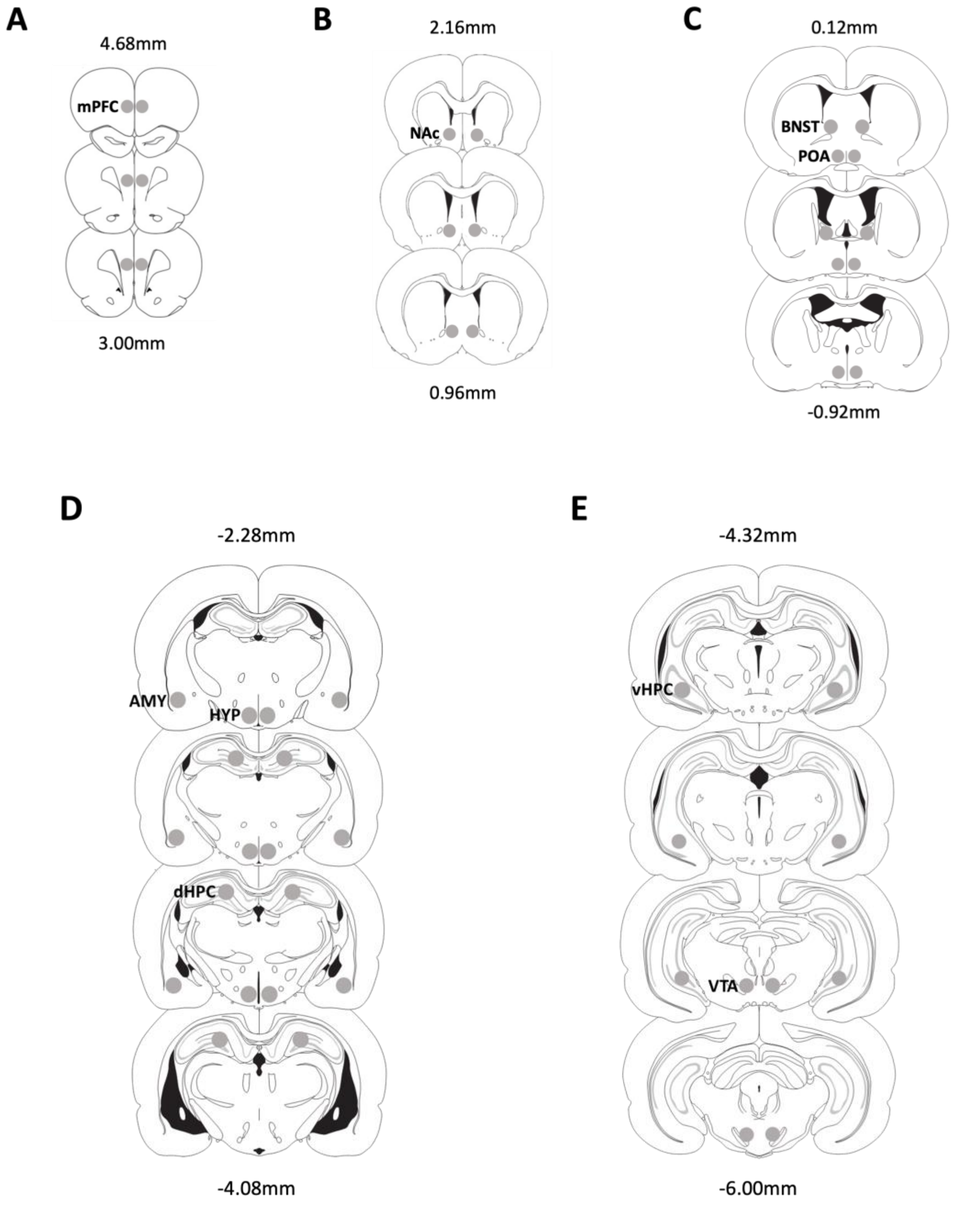


**Figure S1.** Representative bilateral microdissected punch locations taken from fresh frozen tissue sections (300 µm). Coronal brain sections are presented from rostral to caudal; stereotaxic coordinates are presented as proximity to Bregma. 1-mm diameter punches are represented by gray circles. Figure adapted from Tobiansky et al., 2020. Abbreviated brain regions are: medial prefrontal cortex (mPFC), nucleus accumbens (NAc), bed nucleus of the stria terminalis (BNST), preoptic area (POA), hypothalamus (HYP), amygdala (AMY), dorsal hippocampus (dHPC), ventral hippocampus (vHPC), ventral tegmental area (VTA).

**Table 1.** Steroid levels in blood and brain

| | Corticosterone (ng/g) | | DHC (ng/g) | | Progesterone (ng/g) | |
|---|---|---|---|---|---|---|
| | CON | CR | CON | CR | CON | CR |
| Blood | 271.8 ± 18.9 | 288.2 ± 29.1 | 8.3 ± 0.6 | 8.3 ± 0.4 | 17.9 ± 3.5 | 17.1 ± 3.3 |
| Medial prefrontal cortex | 32.1 ± 3.0 | 39.4 ± 3.7 | 3.9 ± 0.4 | 5.0 ± 0.6 | 38.9 ± 6.4 | 43.3 ± 9.0 |
| Nucleus accumbens | 36.6 ± 4.4 | 43.7 ± 3.9 | 4.4 ± 0.5 | 5.1 ± 0.6 | 50.5 ± 9.7 | 47.0 ± 8.9 |
| Bed nucleus of the stria terminalis | 37.0 ± 3.1 | 39.3 ± 4.0 | 5.3 ± 0.6 | 5.0 ± 0.6 | 45.0 ± 7.5 | 40.0 ± 8.1 |
| Preoptic area | 31.7 ± 2.6 | 34.4 ± 3.5 | 6.9 ± 0.9 | 5.9 ± 0.4 | 51.4 ± 8.5 | 45.2 ± 8.8 |
| Hypothalamus | 28.1 ± 3.3 | 32.4 ± 3.2 | 6.6 ± 0.8 | 6.3 ± 0.5 | 39.3 ± 7.0 | 38.7 ± 7.5 |
| Amygdala | 39.5 ± 3.7 | 45.4 ± 4.2 | 4.3 ± 0.5 | 4.2 ± 0.5 | 47.1 ± 8.0 | 46.0 ± 9.4 |
| Dorsal hippocampus | 39.3 ± 3.1 | 44.0 ± 3.7 | 2.6 ± 0.2 | 2.9 ± 0.4 | 42.4 ± 8.4 | 40.4 ± 7.7 |
| Ventral hippocampus | 36.0 ± 2.9 | 44.8 ± 3.9 | 3.5 ± 0.5 | 4.0 ± 0.6 | 41.1 ± 7.0 | 41.3 ± 8.4 |
| Ventral tegmental area | 40.6 ± 3.7 | 51.5 ± 4.8 | 8.2 ± 1.0 | 8.0 ± 0.8 | 64.4 ± 11.4 | 63.7 ± 11.4 |

**Table 2.** Androgens in blood and brain

| | Testosterone (pg/g) | | Androstenedione (pg/g) | |
|---|---|---|---|---|
| | **CON** | **CR** | **CON** | **CR** |
| Blood | 22.1 ± 4.8 | 28.2 ± 5.5 | 149.7 ± 41.8 | 135.2 ± 38.6 |
| Medial prefrontal cortex | 18.1 ± 3.6 | 16.4 ± 5.6 | 167.9 ± 62.1 | 199.2 ± 48.6 |
| Nucleus accumbens | 22.1 ± 7.6 | 30.5 ± 8.4 | 223.9 ± 78.4 | 200.8 ± 44.9 |
| Bed nucleus of the stria terminalis | 16.3 ± 5.5 | 23.3 ± 7.7 | 211.0 ± 72.7 | 194.6 ± 52.8 |
| Preoptic area | 21.4 ± 5.2 | 30.0 ± 8.8 | 246.6 ± 74.3 | 189.0 ± 51.6 |
| Hypothalamus | 19.4 ± 3.4 | 14.3 ± 4.6 | 223.1 ± 82.4 | 182.5 ± 49.7 |
| Amygdala | 23.0 ± 6.9 | 21.1 ± 6.8 | 267.2 ± 84.0 | 229.4 ± 62.9 |
| Dorsal hippocampus | 27.0 ± 10.2 | 23.9 ± 8.5 | 200.0 ± 76.3 | 178.7 ± 45.0 |
| Ventral hippocampus | 24.3 ± 3.4 | 17.5 ± 7.7 | 184.7 ± 60.0 | 209.6 ± 63.0 |
| Ventral tegmental area | 25.2 ± 6.2 | 20.6 ± 11.5 | 307.1 ± 101.0 | 339.4 ± 92.1 |
